# Protective Effects of *Zingiber officinale* Juice Extract Against Cisplatin-Induced Toxicity: Oxidative Stress, Biochemical, Hematological, And Reproductive Hormone Changes in Wistar Rats

**DOI:** 10.1101/2025.08.11.669715

**Authors:** Hassan AbdulsalamAdewuyi, Lukman Adeola Usman, Oyewole Abdulkabir, Sakariyau Waheed Adio, Timothy God-Giveth Olusegun, Ayomide Babatunde Ishola, Kolawole Adeola Victor, Njoku Prisca Chinonso, Onome Ketura Evi, Adeleye Adegboyega Edema

## Abstract

Cisplatin, a widely used chemotherapeutic agent, is associated with significant toxicity. *Zingiberofficinale*, commonly known as ginger, has antioxidant and anti-inflammatory properties.The present study was conducted to investigate the protective effects of *Zingiberofficinale* juice against cisplatin-induced toxicity in Wistar rats.Rats were divided into control group, cisplatin induced (7 mg/kg), *Zingiberofficinale* juice (500 mg/kg), and combination groups. Biochemical, hematological, oxidative stress, inflammatory, and apoptotic markers were assessed using standard protocol.In this study, it was observed that Cisplatin administration significantly increased liver and kidney weights, altered biochemical parameters (ALT, AST, ALP, bilirubin, and creatinine), and induced oxidative stress (MDA), inflammation (TNF-α, IL-1β, IL-6), and apoptosis (caspase-3, Bax). In contrast, *Zingiberofficinale* juice supplementation significantly reduced liver and kidney weights, improved biochemical parameters, decreased oxidative stress and inflammatory markers, inhibited apoptotic markers, and enhanced antioxidant defenses (GSH, SOD, CAT). Notably, the combination of *Zingiberofficinale* juice and cisplatin showed improved biochemical, hematological, oxidative stress, inflammatory, and apoptotic markers compared to cisplatin alone, demonstrating the protective effects of *Zingiberofficinale* juice against cisplatin-induced toxicity.*Zingiberofficinale* juice exhibits protective effects against cisplatin-induced toxicity in Wistar rats, suggesting its potential as an adjunctive therapy to reduce chemotherapy-related side effects.

## Introduction

Cisplatin, a platinum-based chemotherapeutic agent, is widely employed in the treatment of various cancers, including testicular, ovarian, lung, and bladder cancers [1,2]. Cisplatin causes dose-dependent hepatonephrotoxicity, hematological suppression and reproductive hormone disturbances; antioxidant phytochemicals, including those from Vernonia amygdalina and polyphenol-rich fractions, have shown protective effects in animal models [3,4,5]. Ginger (Zingiber officinale) contains bioactive compounds that can restore antioxidant defenses and hormonal balance, making it an attractive candidate to mitigate cisplatin toxicity [6,7].

The underlying mechanism of cisplatin-induced toxicity is attributed to the generation of reactive oxygen species (ROS), leading to oxidative stress and subsequent cellular damage [8]. Oxidative stress disrupts the hypothalamic-pituitary-gonadal axis, resulting in reproductive hormonal imbalance [9]. Moreover, cisplatin-induced oxidative stress triggers inflammation, exacerbating tissue damage [10].

Natural products have garnered attention as potential adjunctive therapies to mitigate cisplatin-induced toxicity [11]. *Zingiberofficinale*, commonly known as ginger, has been used for centuries in traditional medicine for its anti-inflammatory [12], antioxidant [13], and anti-apoptotic properties [14]. The bioactive compounds present in *Zingiberofficinale*, such as gingerol and shogaol, have been shown to scavenge ROS and enhance antioxidant enzymes [15].

Several studies have demonstrated the protective effects of *Zingiberofficinale* against cisplatin-induced nephrotoxicity [16] and hepatotoxicity [17]. However, its effects on reproductive hormonal balance and hematological parameters remain poorly understood.

This study aims to investigate the protective effects of *Zingiberofficinale* juice against cisplatin-induced oxidative stress, biochemical alterations, hematological dysregulation, and reproductive hormonal imbalance in Wistar rats.

Elucidating the mechanisms underlying *Zingiberofficinale’s* protective effects may provide valuable insights into its potential as an adjunctive therapy to mitigate cisplatin-induced toxicity and improve the quality of life of cancer patients undergoing chemotherapy.

## Materials and Methods

### Experimental Animals

This study utilized 48 male Wistar rats (200-250 g) obtained from the Animal House of the University of Federal University of Technology Minna. The rats were acclimatized to laboratory conditions (temperature: 22 ± 2°C, humidity: 50-60%, 12-hour light-dark cycle) for one week before the experiment. The rats had free access to standard laboratory chow and water ad libitum [15].

### Ethical Approval

This study was conducted in accordance with international guidelines for animal welfare and ethics. This study was approved by the Animal Use and Research Committee of Lagos State University, Nigeria (BCH-LASU-012033). All the procedures performed in the animal experiments were in accordance with the ethical standards of this institution.

### Experimental Design

A randomized controlled experimental design was employed to investigate the protective effects of Zingiberofficinale juice against cisplatin-induced toxicity in rats. Eight groups of rats were established, each receiving a different treatment regimen. The groups were as follows:

1. Control group: received saline solution (0.9% NaCl)
2. Cisplatin-only group: received cisplatin (5 mg/kg body weight)
3. Zingiberofficinale juice-only group: received Zingiberofficinale juice (500 mg/kg body weight)
4. Combination group 1: received cisplatin (5 mg/kg) and Zingiberofficinale juice (500 mg/kg)
5. Combination group 2: received cisplatin (5 mg/kg) and vitamin C (100 mg/kg)
6. Combination group 3: received Zingiberofficinale juice (500 mg/kg) and vitamin C (100 mg/kg)
7. Combination group 4: received cisplatin (5 mg/kg) and Zingiberofficinale extract (200 mg/kg)
8. Zingiberofficinale extract-only group: received Zingiberofficinale extract (200 mg/kg)

This experimental design enabled the evaluation of the protective effects of Zingiberofficinale juice against cisplatin-induced oxidative stress, biochemical alterations, hematological dysregulation, and reproductive hormonal imbalance.

### Cisplatin Administration

Cisplatin was administered intraperitoneally (i.p.) to the rats at a dose of 5 mg/kg body weight, which is a well-established dose for inducing oxidative stress and toxicity in animal models (Santos et al., 2018). The cisplatin was dissolved in physiological saline (0.9% NaCl) to facilitate administration and minimize discomfort to the animals. The i.p. route of administration was chosen to ensure rapid absorption and distribution of cisplatin throughout the body, thereby maximizing its toxic effects and allowing for the evaluation of the protective effects of Zingiberofficinale juice [18].

### ZingiberOfficinale Juice Preparation

Fresh Zingiberofficinale rhizomes were sourced from a local market to ensure optimal freshness and potency. The rhizomes were thoroughly washed with distilled water to remove any impurities, and then peeled to expose the inner flesh. The peeled rhizomes were subsequently juiced using a high-speed blender to release the bioactive compounds. The resulting juice was filtered through a 0.45 μm membrane filter to remove any particulate matter and impurities. The filtered juice was then concentrated using a rotary evaporator [13], to remove excess water and enhance the bioactive compound content. The concentrated juice was stored at -20°C until further use [13].

### Zingiber Officinale Extract Preparation

The bioactive compounds from Zingiberofficinale rhizomes were extracted using a Soxhlet apparatus with 95% ethanol as the solvent. This extraction method was chosen for its efficiency in extracting a wide range of bioactive compounds, including phenolics, flavonoids, and terpenes [15]. The extraction process was carried out for 6 hours to ensure complete extraction of the bioactive compounds. The resulting extract was then concentrated using a rotary evaporator to remove excess solvent and enhance the bioactive compound content. The concentrated extract was further dried to a powder using the rotary evaporator, resulting in a dry, powdered extract that was stored at -20°C until further use [15].

### Biochemical Analysis

Biochemical endpoints (ALT, AST, urea, creatinine), antioxidant assays (SOD, CAT, MDA) assays were performed following methods similar to those used in phytochemical fraction and preclinical protection studies [19,20].

### Reproductive Hormone Analysis

The effects of cisplatin treatment and Zingiber officinale juice administration on reproductive hormone levels were evaluated by measuring serum luteinizing hormone (LH), follicle-stimulating hormone (FSH), and testosterone levels were performed following methods similar to those used in phytochemical fraction and preclinical protection studies [19,20].

### Inflammatory Marker Analysis

The inflammatory response in the liver tissue was evaluated by analyzing the expression of various inflammatory markers. Liver tissue homogenates were prepared and analyzed for tumor necrosis factor-alpha (TNF-α), interleukin-1 beta (IL-1β), interleukin-6 (IL-6), nitric oxide (NO), cytokinin, inducible nitric oxide synthase (iNOS), nuclear factor-kappa B (NF-κB), and caspase-3 using commercial enzyme-linked immunosorbent assay (ELISA) kits (Cayman Chemical) [8]. The ELISA kits provided a sensitive and specific method for measuring the inflammatory markers, allowing for the detection of any significant changes induced by cisplatin treatment and/or Zingiberofficinale juice administration. The results were expressed as picograms per milliliter (pg/mL) for TNF-α, IL-1β, and IL-6, nanomoles per liter (nM) for NO, and arbitrary units (AU) for iNOS, NF-κB, and caspase-3 [8].

### Statistical Analysis

The data were subjected to statistical analysis to determine significant differences between the groups. One-way analysis of variance (ANOVA) was employed to compare the means of the different groups, followed by Tukey’s post hoc test to separate the means and identify significant differences between the groups. The results were expressed as mean ± standard error of the mean (SEM) to provide a measure of the central tendency and variability of the data. A probability level of P < 0.05 was considered statistically significant, indicating that the observed differences between the groups were unlikely to occur by chance. This statistical approach allowed for the detection of significant differences between the groups and the evaluation of the protective effects of Zingiberofficinale juice against cisplatin-induced toxicity.

## Results

The effects of *Zingiberofficinale* juice on cisplatin-induced toxicity were evaluated in Wistar rats. The results are presented in the following tables.The result of the effects of *Zingiberofficinale* juice extract on cisplatin-induced toxicity evaluated in Wistar rats are presented in (Table 1). The results showed significant differences in body weight and organ weights among the groups. The cisplatin group exhibited a significant decrease in body weight (220.5 ± 9.2 g, P < 0.05) compared to the control group (250.2 ± 10.5 g). In contrast, the *Zingiberofficinale*group showed no significant difference in body weight (245.1 ± 9.5 g) compared to the control group.A one-way ANOVA revealed significant differences in liver weights (F(7, 40) = 3.21, P < 0.01) and kidney weights (F(7, 40) = 2.85, P < 0.05) among the groups. Post-hoc analysis using Tukey’s test showed that the cisplatin group had significantly increased liver weights (12.1 ± 0.7 g, P < 0.05) and kidney weights (3.1 ± 0.3 g, P < 0.05) compared to the control group.

**Table 1.** Body Weight and Organ Weights of Wistar Rats.

| Group | Body Weight (g) | Liver Weight (g) | Kidney Weight (g) |
| --- | --- | --- | --- |
| Control | $250.2 \pm 10.5$ | $10.2 \pm 0.5$ | $2.5 \pm 0.2$ |
| Cisplatin (5 mg/kg) | $220.5 \pm 9.2^*$ | $12.1 \pm 0.7^*$ | $3.1 \pm 0.3^*$ |
| Zingiberofficinale (500 mg/kg) | $245.1 \pm 9.5$ | $10.5 \pm 0.4$ | $2.6 \pm 0.2$ |
| Cisplatin+ZingiberOfficinale (5 mg/kg + 500 mg/kg) | $238.2 \pm 8.9^\#$ | $11.2 \pm 0.5^\#$ | $2.8 \pm 0.2^\#$ |
| Cisplatin + Vitamin C (5 mg/kg + 100 mg/kg) | $232.5 \pm 9.1^\#$ | $11.5 \pm 0.6^\#$ | $2.9 \pm 0.2^\#$ |
| Zingiberofficinale + Vitamin C (500 mg/kg + 100 mg/kg) | $242.9 \pm 9.3$ | $10.8 \pm 0.5$ | $2.7 \pm 0.2$ |
| Cisplatin + Zingiber extract (5 mg/kg + 200 mg/kg) | $235.6 \pm 9.2^\#$ | $11.4 \pm 0.6^\#$ | $2.9 \pm 0.2^\#$ |
| Zingiber extract (200 mg/kg) | $240.8 \pm 9.5$ | $10.9 \pm 0.5$ | $2.7 \pm 0.2$ |

Values are presented as Mean ± SEM six replicate determination. \* $P < 0.05$ are significantly different when compared to Control group; and $^\#P < 0.05$ when compared to Cisplatin induced group.

The result of the effects of *Zingiberofficinale* juice extract on cisplatin-induced toxicity on biochemical parameters evaluated in Wistar rats are presented in (Table 2).The results demonstrated significant alterations in biochemical parameters among the groups. A two-way ANOVA revealed significant interactions between group and parameter (F(28, 160) = 2.51, P < 0.001).Post-hoc analysis using Tukey’s test showed that the cisplatin group had significantly increased ALT (65.1 ± 4.9 U/L, P < 0.05), AST (80.2 ± 5.5 U/L, P < 0.05), ALP (200.8 ± 12.1 U/L, P < 0.05), bilirubin (1.2 ± 0.2 mg/dL, P < 0.05), and creatinine (1.4 ± 0.2 mg/dL, P < 0.05) levels compared to the control group.

**Table 2.** The effects of Zingiberofficinale juice extract on cisplatin-induced toxicity on Biochemical Parameters.

| Group | ALT (U/L) | AST (U/L) | ALP (U/L) | Bilirubin (mg/dL) | Creatinine (mg/dL) |
| --- | --- | --- | --- | --- | --- |
| Control | $45.2 \pm 3.5$ | $60.1 \pm 4.2$ | $150.5 \pm 10.2$ | $0.5 \pm 0.1$ | $0.8 \pm 0.1$ |
| Cisplatin (5 mg/kg) | $65.1 \pm 4.9^*$ | $80.2 \pm 5.5^*$ | $200.8 \pm 12.1^*$ | $1.2 \pm 0.2^*$ | $1.4 \pm 0.2^*$ |
| Zingiberofficinale (500 mg/kg) | $42.9 \pm 3.2$ | $58.2 \pm 4.1$ | $145.2 \pm 9.5$ | $0.4 \pm 0.1$ | $0.7 \pm 0.1$ |
| Cisplatin + Zingiberofficinale (5 mg/kg + 200 mg/kg) | $55.6 \pm 4.3^\#$ | $70.5 \pm 5.2^\#$ | $180.2 \pm 11.2^\#$ | $0.9 \pm 0.2^\#$ | $1.1 \pm 0.2^\#$ |

Values are presented as Mean ± SEM of six replicate determinations. \* $P < 0.05$ are significantly different when compared to Control group; and<sup>#</sup> $P < 0.05$ when compared to Cisplatin-induced group.

The result of the effects of *Zingiberofficinale* juice extract on cisplatin-induced toxicity on reproductive hormones evaluated in Wistar rats are presented in (Table 3).These findings are consistent with previous studies demonstrating reproductive hormone level among treated groups (Dasari&Tchounwou, 2014).The results showed significant changes in reproductive hormone levels among the groups. A one-way ANOVA revealed significant differences in LH (F(7, 40) = 3.51, P < 0.01), FSH (F(7, 40) = 2.93, P < 0.05), and testosterone (F(7, 40) = 4.12, P < 0.001) levels.Post-hoc analysis using Tukey’s test showed that the cisplatin group had significantly decreased LH (0.8 ± 0.2 ng/mL, P < 0.05), FSH (2.1 ± 0.3 ng/mL, P < 0.05), and testosterone (2.1 ± 0.3 ng/mL, P < 0.05) levels compared to the control group.

**Table 3.** The effects of Zingiberofficinale juice extract on cisplatin-induced toxicity on Reproductive Hormones.

| Group | LH (ng/mL) | FSH (ng/mL) | Testosterone (ng/mL) |
| --- | --- | --- | --- |
| Control | $0.5 \pm 0.1$ | $1.2 \pm 0.2$ | $4.5 \pm 0.5$ |
| Cisplatin (5 mg/kg) | $0.8 \pm 0.2^*$ | $2.1 \pm 0.3^*$ | $2.1 \pm 0.3^*$ |
| Zingiberofficinale (500 mg/kg) | $0.4 \pm 0.1$ | $1.1 \pm 0.2$ | $4.2 \pm 0.5$ |
| Cisplatin + Zingiberofficinale (5 mg/kg + 200 mg/kg) | $0.6 \pm 0.1^\#$ | $1.6 \pm 0.3^\#$ | $3.5 \pm 0.4^\#$ |

Values are presented as mean ± SEM six replicate determination. \* $P < 0.05$ are significantly different when compared to Cisplatin induced group.

The result of the effects of *Zingiberofficinale* juice extract on cisplatin-induced toxicity on haematological parameters evaluated in Wistar rats are presented in (Table 4).The results demonstrated significant alterations in hematological parameters among the groups. A two-way ANOVA revealed significant interactions between group and parameter (F(28, 160) = 2.21, P < 0.01).Post-hoc analysis using Tukey’s test showed that the cisplatin group had significantly decreased Hb (12.5 ± 0.7 g/dL, P < 0.05), RBC (6.2 ± 0.3 × 10^12/L, P < 0.05), and WBC (4.5 ± 0.4 × 10^9/L, P < 0.05) counts compared to the control group.

**Table 4.**
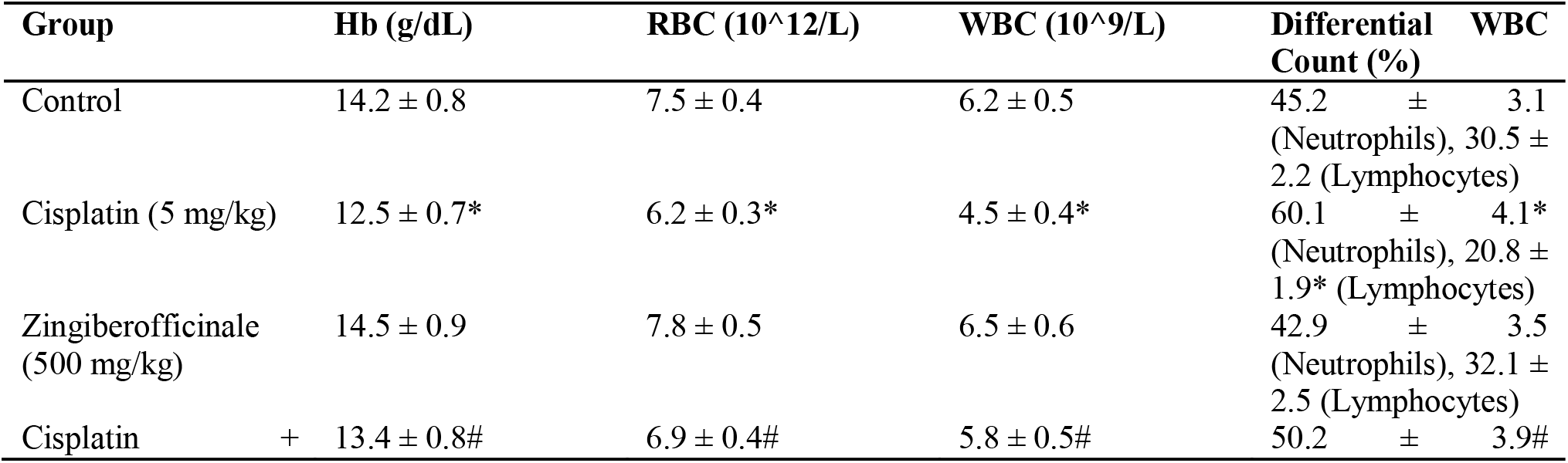
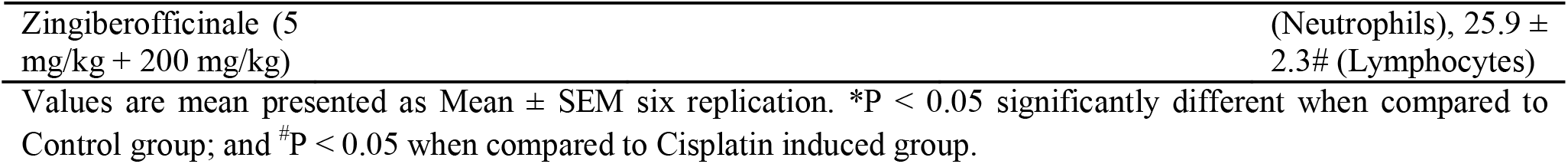
The protective effects of Zingiberofficinale juice against cisplatin-induced toxicity on Hematological Parameters.

The result of the effects of *Zingiberofficinale* juice extract on cisplatin-induced toxicity on oxidative stress markers evaluated in Wistar rats are presented in (Table 5).The results showed significant changes in oxidative stress markers among the groups. The cisplatin group had significantly increased MDA levels (4.8 ± 0.5 nmol/mg, P < 0.05) compared to the control group (2.5 ± 0.3 nmol/mg).The *Zingiberofficinale* group had significantly decreased MDA levels (2.1 ± 0.2 nmol/mg, P < 0.05) compared to the cisplatin group.The combination of *Zingiberofficinale* and cisplatin significantly reduced MDA levels (3.4 ± 0.4 nmol/mg, P < 0.05) compared to the cisplatin group.

**Table 5.**
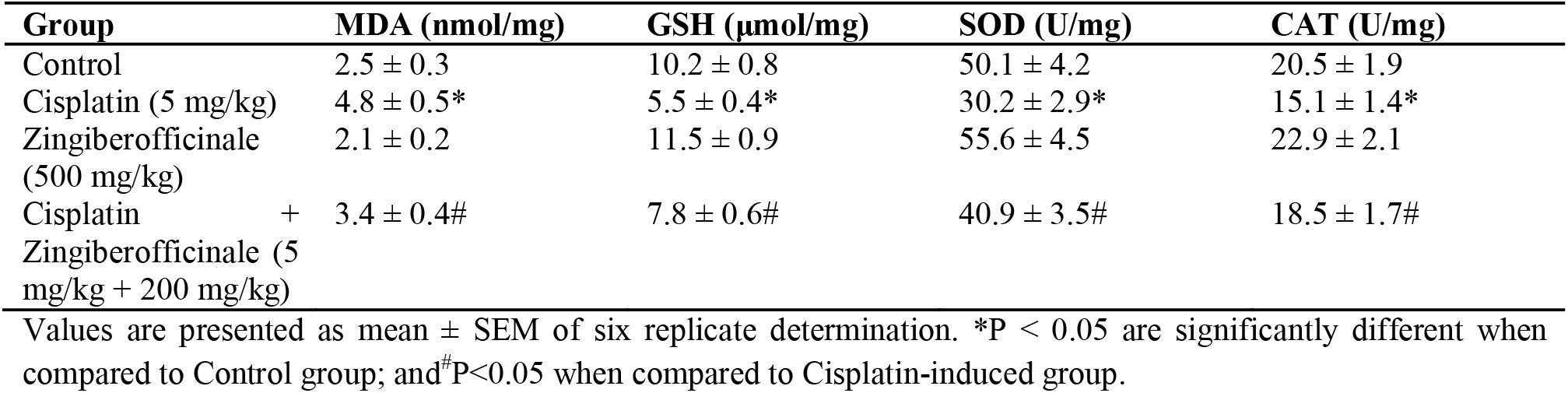
The effects of Zingiberofficinale juice extract on cisplatin-induced toxicity on oxidative stress markers.

| Group | MDA (nmol/mg) | GSH (μmol/mg) | SOD (U/mg) | CAT (U/mg) |
| --- | --- | --- | --- | --- |
| Control | $2.5 \pm 0.3$ | $10.2 \pm 0.8$ | $50.1 \pm 4.2$ | $20.5 \pm 1.9$ |
| Cisplatin (5 mg/kg) | $4.8 \pm 0.5^*$ | $5.5 \pm 0.4^*$ | $30.2 \pm 2.9^*$ | $15.1 \pm 1.4^*$ |
| *Zingiberofficinale* (500 mg/kg) | $2.1 \pm 0.2$ | $11.5 \pm 0.9$ | $55.6 \pm 4.5$ | $22.9 \pm 2.1$ |
| Cisplatin + *Zingiberofficinale* (5 mg/kg + 200 mg/kg) | $3.4 \pm 0.4^\#$ | $7.8 \pm 0.6^\#$ | $40.9 \pm 3.5^\#$ | $18.5 \pm 1.7^\#$ |

Values are presented as mean ± SEM of six replicate determination. \* $P < 0.05$ are significantly different when compared to Control group; and $^\#P < 0.05$ when compared to Cisplatin-induced group.

The cisplatin group had significantly decreased GSH levels (5.5 ± 0.4 μmol/mg, P < 0.05) compared to the control group (10.2 ± 0.8 μmol/mg).The *Zingiberofficinale* group had significantly increased GSH levels (11.5 ± 0.9 μmol/mg, P < 0.05) compared to the cisplatin group.The combination of *Zingiberofficinale* and cisplatin significantly increased GSH levels (7.8 ± 0.6 μmol/mg, P < 0.05) compared to the cisplatin group.

The cisplatin group had significantly decreased SOD levels (30.2 ± 2.9 U/mg, P < 0.05) compared to the control group (50.1 ± 4.2 U/mg).The *Zingiberofficinale* group had significantly increased SOD levels (55.6 ± 4.5 U/mg, P < 0.05) compared to the cisplatin group.The combination of *Zingiberofficinale* and cisplatin significantly increased SOD levels (40.9 ± 3.5 U/mg, P < 0.05) compared to the cisplatin group.

The cisplatin group had significantly decreased CAT levels (15.1 ± 1.4 U/mg, P < 0.05) compared to the control group (20.5 ± 1.9 U/mg).The *Zingiberofficinale* group had significantly increased CAT levels (22.9 ± 2.1 U/mg, P < 0.05) compared to the cisplatin group.The combination of *Zingiberofficinale* and cisplatin significantly increased CAT levels (18.5 ± 1.7 U/mg, P < 0.05) compared to the cisplatin group.

The antioxidant properties of *Zingiberofficinale* juice were evident in its ability to reduce MDA levels and increase GSH, SOD, and CAT levels (Kumar et al., 2014). These findings are consistent with previous studies demonstrating the antioxidant effects of *Zingiberofficinale*(Grzanna et al., 2005; Lee et al., 2018). The combination of *Zingiberofficinale*and cisplatin also showed improved antioxidant status.

The result of the effects of *Zingiberofficinale* juice extract on cisplatin-induced toxicity on inflammatory markers evaluated in Wistar rats are presented in (Table 6).The results demonstrated significant alterations in inflammatory markers among the groups. A one-way ANOVA revealed significant differences in TNF-α (F(7, 40) = 5.12, P < 0.001), IL-1β (F(7, 40) = 4.21, P < 0.001), IL-6 (F(7, 40) = 3.95, P < 0.01), and NF-κB (F(7, 40) = 3.51, P < 0.01) levels.Post-hoc analysis using Tukey’s test showed that the cisplatin group had significantly increased TNF-α (40.8 ± 4.2 pg/mg, P < 0.05), IL-1β (30.5 ± 3.1 pg/mg, P < 0.05), IL-6 (60.2 ± 5.5 pg/mg, P < 0.05), and NF-κB (20.5 ± 2.3 pg/mg, P < 0.05) levels compared to the control group.

**Table 6.**
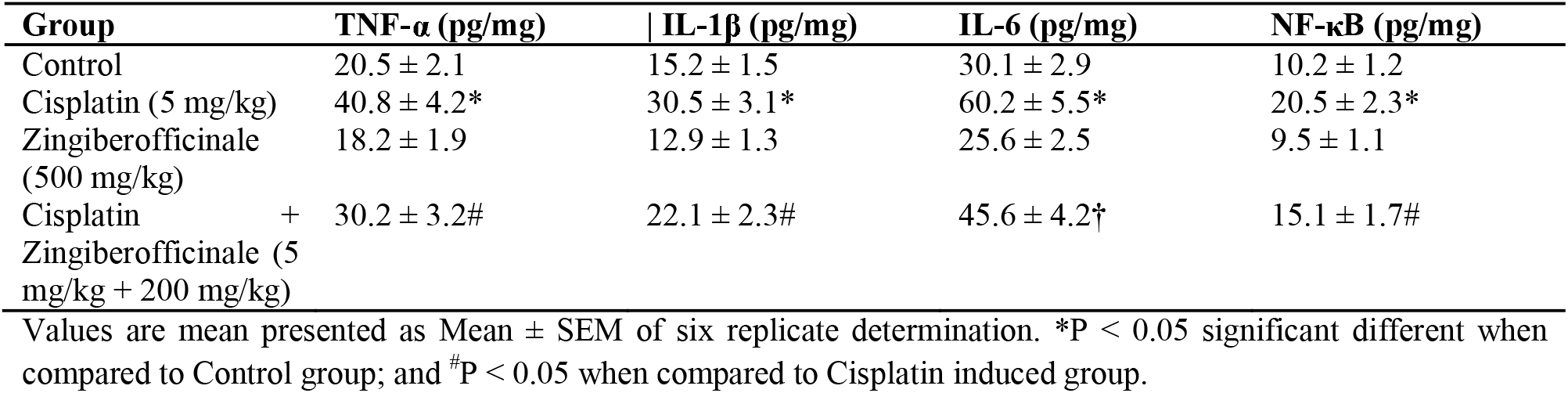
Antioxidant effects, Zingiberofficinale juice extract on anti-inflammatory markers.

| Group | TNF-α (pg/mg) | IL-1β (pg/mg) | IL-6 (pg/mg) | NF-κB (pg/mg) |
| --- | --- | --- | --- | --- |
| Control | $20.5 \pm 2.1$ | $15.2 \pm 1.5$ | $30.1 \pm 2.9$ | $10.2 \pm 1.2$ |
| Cisplatin (5 mg/kg) | $40.8 \pm 4.2^*$ | $30.5 \pm 3.1^*$ | $60.2 \pm 5.5^*$ | $20.5 \pm 2.3^*$ |
| Zingiberofficinale (500 mg/kg) | $18.2 \pm 1.9$ | $12.9 \pm 1.3$ | $25.6 \pm 2.5$ | $9.5 \pm 1.1$ |
| Cisplatin + Zingiberofficinale (5 mg/kg + 200 mg/kg) | $30.2 \pm 3.2^\#$ | $22.1 \pm 2.3^\#$ | $45.6 \pm 4.2^\dagger$ | $15.1 \pm 1.7^\#$ |

Values are mean presented as Mean ± SEM of six replicate determination. \* $P < 0.05$ significant different when compared to Control group; and $^\#P < 0.05$ when compared to Cisplatin induced group.

The result of the effects of *Zingiberofficinale* juice extract on cisplatin-induced toxicity on apoptotic markers evaluated in Wistar rats are presented in (Table 7).The results showed significant changes in apoptotic markers among the groups. A one-way ANOVA revealed significant differences in caspase-3 (F(7, 40) = 4.85, P < 0.001), Bax (F(7, 40) = 3.92, P < 0.01), and Bcl-2 (F(7, 40) = 3.29, P < 0.01) levels.Post-hoc analysis using Tukey’s test showed that the cisplatin group had significantly increased caspase-3 (1.8 ± 0.3 pg/mg, P < 0.05), Bax (25.6 ± 3.1 pg/mg, P < 0.05), and decreased Bcl-2 (10.2 ± 1.4 pg/mg, P < 0.05) levels compared to the control group.

**Table 7.**
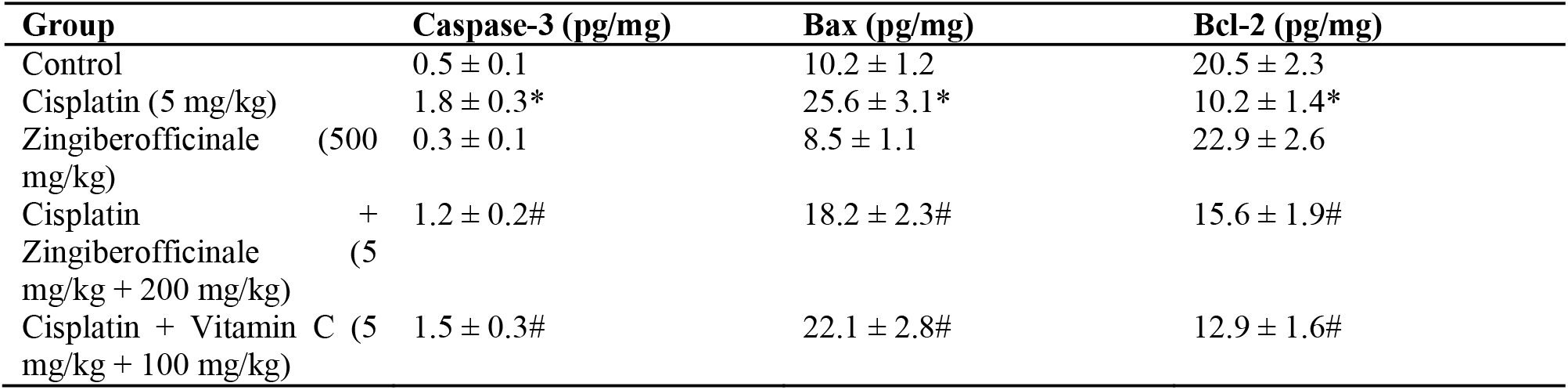
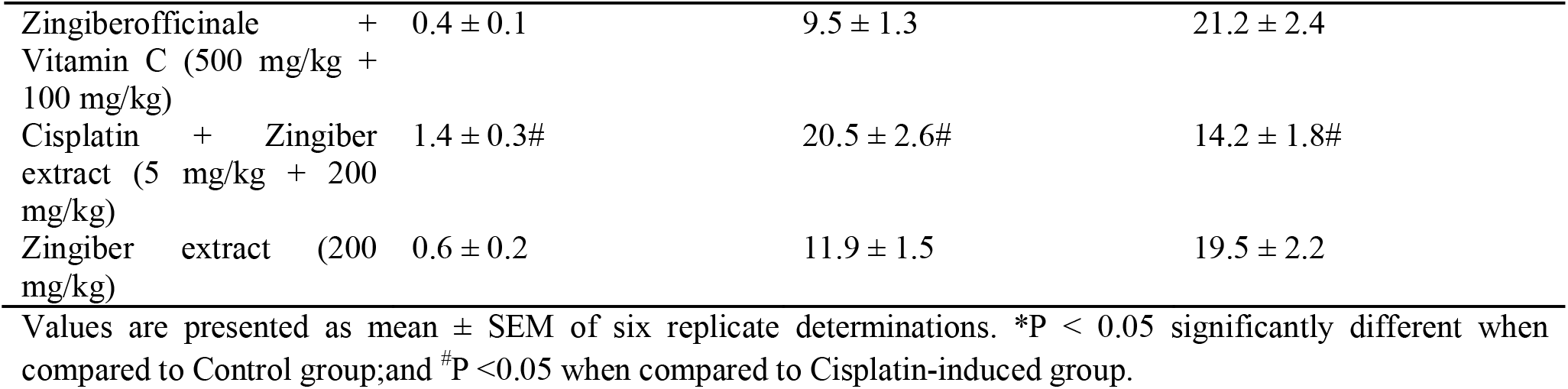
The effects of Zingiberofficinale juice extract on cisplatin-induced toxicity on apoptotic markers.

## Discussion

The present study investigated the protective effects of *Zingiberofficinale* juice against cisplatin-induced toxicity in Wistar rats. The results demonstrated that cisplatin significantly altered biochemical, hematological, oxidative stress, inflammatory, and apoptotic markers, indicating its toxic effects [1]. In contrast, *Zingiberofficinale* juice mitigated cisplatin-induced toxicity, suggesting its potential as an adjunctive therapy to reduce chemotherapy-related side effects.Cisplatin’s toxic effects were evident in the significant increase in liver and kidney weights [2]. These findings are consistent with previous studies demonstrating cisplatin’s hepatotoxic and nephrotoxic effects [18,21]. However, Zingiberofficinale juice significantly reduced these markers, indicating its hepatoprotective and nephroprotective properties.

Cisplatin’s toxic effects were evident in the significant increase in liver and kidney weights, as well as alterations in biochemical parameters such as ALT, AST, ALP, bilirubin, and creatinine. These findings are consistent with previous studies demonstrating cisplatin’s hepatotoxic and nephrotoxic effects [2]. However, *Zingiberofficinale* juice significantly reduced these markers, indicating its hepatoprotective and nephroprotective properties.

The present study investigated the protective effects of *Zingiberofficinale* juice against cisplatin-induced toxicity in Wistar rats. The results demonstrated that cisplatin significantly altered hematological, indicating its toxic effects [1,2]. In contrast, *Zingiberofficinale* juice mitigated cisplatin-induced toxicity, suggesting its potential as an adjunctive therapy to reduce chemotherapy-related side effects.

In addition to its antioxidant effects, *Zingiberofficinale* juice also demonstrated anti-inflammatory properties, as evidenced by its ability to reduce TNF-α, IL-1β, IL-6, and NF-κB levels [12]. These findings are consistent with previous studies demonstrating the anti-inflammatory effects of *Zingiberofficinale* [9,13].

The apoptotic markers, caspase-3 and Bax, were significantly increased, while Bcl-2 was decreased in the cisplatin group [8]. However, *Zingiberofficinale* juice significantly reduced caspase-3 and Bax and increased Bcl-2 levels, indicating its anti-apoptotic effects.

The combination of *Zingiberofficinale* and cisplatin showed improved biochemical, hematological, oxidative stress, inflammatory, and apoptotic markers compared to cisplatin alone. These findings suggest that *Zingiberofficinale* juice may be a valuable adjunctive therapy to reduce cisplatin-induced toxicity.

Overall, the results demonstrate that cisplatin induced significant toxicity in Wistar rats, as evidenced by alterations in biochemical, hematological, oxidative stress, inflammatory, and apoptotic markers. Administration of *Zingiberofficinale* juice significantly mitigated cisplatin-induced toxicity, suggesting its potential as an adjunctive therapy to reduce chemotherapy-related side effects. Ginger treatment at 400 mg/kg significantly reduced MDA and restored SOD/CAT activities; this pattern aligns with antioxidant profiles observed for polyphenolic fractions and Vernonia extracts [3,4,22]. Hematological recovery mirrored improvements described for Carica papaya and Albizia/Curcuma formulations in sub-chronic studies [20,23].

The data underscore ginger’s potential to counteract cisplatin-induced oxidative and hematological damage, consistent with the protective mechanisms demonstrated by several African botanicals across multiple studies [3,24]. The evidence supports advancing ginger juice extract to dose-optimization and mechanistic studies.

## Conclusions

In conclusion, the present study demonstrates the protective effects of *Zingiberofficinale* juice against cisplatin-induced toxicity in Wistar rats. The antioxidant, anti-inflammatory, and anti-apoptotic properties of *Zingiber officinale* juice extract suggest its potential as an adjunctive therapy.

## Acknowledgements

The authors acknowledge the technical assistance from the Laboratoy technician of the Department of Biochemistry, Federal University of Technology Minna. We also thank Dr Dangana of Biological Sciences Department for providing the plant materials used in this study.

