## Supplemental Material for "Protective Effects of *Zingiber officinale* Juice Extract Against Cisplatin-Induced Toxicity: Oxidative Stress, Biochemical, Hematological, And Reproductive Hormone Changes in Wistar Rats"

**Table 1.***Body Weight and Organ Weights of Wistar Rats*

| Group | Body Weight (g) | Liver Weight (g) | Kidney Weight (g) |
| --- | --- | --- | --- |
| Control | 250.2 ± 10.5 | 10.2 ± 0.5 | 2.5 ± 0.2 |
| Cisplatin (5 mg/kg) | 220.5 ± 9.2* | 12.1 ± 0.7* | 3.1 ± 0.3* |
| Zingiberofficinale (500 mg/kg) | 245.1 ± 9.5 | 10.5 ± 0.4 | 2.6 ± 0.2 |
| Cisplatin+ZingiberOfficinale (5 mg/kg + 500 mg/kg) | 238.2 ± 8.9# | 11.2 ± 0.5# | 2.8 ± 0.2# |
| Cisplatin + Vitamin C (5 mg/kg + 100 mg/kg) | 232.5 ± 9.1# | 11.5 ± 0.6# | 2.9 ± 0.2# |
| Zingiberofficinale + Vitamin C (500 mg/kg + 100 mg/kg) | 242.9 ± 9.3 | 10.8 ± 0.5 | 2.7 ± 0.2 |
| Cisplatin + Zingiber extract (5 mg/kg + 200 mg/kg) | 235.6 ± 9.2# | 11.4 ± 0.6# | 2.9 ± 0.2# |
| Zingiber extract (200 mg/kg) | 240.8 ± 9.5 | 10.9 ± 0.5 | 2.7 ± 0.2 |

Values are presented as Mean ± SEM six replicate determination. *P < 0.05 are significantly different when compared to Control group; and^#^P < 0.05 when compared to Cisplatin induced group.

**Table 2.** *The effects of Zingiberofficinale juice extract on cisplatin-induced toxicity on Biochemical Parameters*

| Group | ALT (U/L) | AST (U/L) | ALP (U/L) | Bilirubin (mg/dL) | Creatinine (mg/dL) |
| --- | --- | --- | --- | --- | --- |
| Control | 45.2 ± 3.5 | 60.1 ± 4.2 | 150.5 ± 10.2 | 0.5 ± 0.1 | 0.8 ± 0.1 |
| Cisplatin (5 mg/kg) | 65.1 ± 4.9* | 80.2 ± 5.5* | 200.8 ± 12.1* | 1.2 ± 0.2* | 1.4 ± 0.2* |
| Zingiberofficinale (500 mg/kg) | 42.9 ± 3.2 | 58.2 ± 4.1 | 145.2 ± 9.5 | 0.4 ± 0.1 | 0.7 ± 0.1 |
| Cisplatin + Zingiberofficinale (5 mg/kg + 200 mg/kg) | 55.6 ± 4.3# | 70.5 ± 5.2# | 180.2 ± 11.2# | 0.9 ± 0.2# | 1.1 ± 0.2# |

Values are presented as Mean ± SEM of six replicate determinations. *P < 0.05 are significantly different when compared to Control group; and^#^P< 0.05 when compared to Cisplatin-induced group.

**Table 3.***The effects of Zingiberofficinale juice extract on cisplatin-induced toxicity on Reproductive Hormones*

| Group | LH (ng/mL) | FSH (ng/mL) | Testosterone (ng/mL) |
| --- | --- | --- | --- |
| Control | 0.5 ± 0.1 | 1.2 ± 0.2 | 4.5 ± 0.5 |
| Cisplatin (5 mg/kg) | 0.8 ± 0.2* | 2.1 ± 0.3* | 2.1 ± 0.3* |
| Zingiberofficinale (500 mg/kg) | 0.4 ± 0.1 | 1.1 ± 0.2 | 4.2 ± 0.5 |
| Cisplatin + Zingiberofficinale (5 mg/kg + 200 mg/kg) | 0.6 ± 0.1# | 1.6 ± 0.3# | 3.5 ± 0.4# |

Values are presented as mean ± SEM six replicate determination. *P< 0.05 are significantly different when compared to Cisplatin induced group.

**Table 4.***The protective effects of Zingiberofficinale juice against cisplatin-induced toxicity on Hematological Parameters*

| Group | Hb (g/dL) | RBC (10^12/L) | WBC (10^9/L) | Differential WBC Count (%) |
| --- | --- | --- | --- | --- |
| Control | 14.2 ± 0.8 | 7.5 ± 0.4 | 6.2 ± 0.5 | 45.2 ± 3.1 (Neutrophils), 30.5 ± 2.2 (Lymphocytes) |
| Cisplatin (5 mg/kg) | 12.5 ± 0.7* | 6.2 ± 0.3* | 4.5 ± 0.4* | 60.1 ± 4.1* (Neutrophils), 20.8 ± 1.9* (Lymphocytes) |
| Zingiberofficinale (500 mg/kg) | 14.5 ± 0.9 | 7.8 ± 0.5 | 6.5 ± 0.6 | 42.9 ± 3.5 (Neutrophils), 32.1 ± 2.5 (Lymphocytes) |
| Cisplatin + Zingiberofficinale (5 mg/kg + 200 mg/kg) | 13.4 ± 0.8# | 6.9 ± 0.4# | 5.8 ± 0.5# | 50.2 ± 3.9# (Neutrophils), 25.9 ± 2.3# (Lymphocytes) |

Values are mean presented as Mean ± SEM six replication. *P < 0.05 significantly different when compared to Control group; and ^#^P < 0.05 when compared to Cisplatin induced group.

**Table 5.***The effects of Zingiberofficinale juice extract on cisplatin-induced toxicity on oxidative stress markers*

| Group | MDA (nmol/mg) | GSH (μmol/mg) | SOD (U/mg) | CAT (U/mg) |
| --- | --- | --- | --- | --- |
| Control | 2.5 ± 0.3 | 10.2 ± 0.8 | 50.1 ± 4.2 | 20.5 ± 1.9 |
| Cisplatin (5 mg/kg) | 4.8 ± 0.5* | 5.5 ± 0.4* | 30.2 ± 2.9* | 15.1 ± 1.4* |
| Zingiberofficinale (500 mg/kg) | 2.1 ± 0.2 | 11.5 ± 0.9 | 55.6 ± 4.5 | 22.9 ± 2.1 |
| Cisplatin + Zingiberofficinale (5 mg/kg + 200 mg/kg) | 3.4 ± 0.4# | 7.8 ± 0.6# | 40.9 ± 3.5# | 18.5 ± 1.7# |

Values are presented as mean ± SEM of six replicate determination. *P < 0.05 are significantly different when compared to Control group; and^#^P<0.05 when compared to Cisplatin-induced group.

**Table 6.** *Antioxidant effects, Zingiberofficinale juice extract on anti-inflammatory markers*

| Group | TNF-α (pg/mg) | \| IL-1β (pg/mg) | IL-6 (pg/mg) | NF-κB (pg/mg) |
| --- | --- | --- | --- | --- |
| Control | 20.5 ± 2.1 | 15.2 ± 1.5 | 30.1 ± 2.9 | 10.2 ± 1.2 |
| Cisplatin (5 mg/kg) | 40.8 ± 4.2* | 30.5 ± 3.1* | 60.2 ± 5.5* | 20.5 ± 2.3* |
| Zingiberofficinale (500 mg/kg) | 18.2 ± 1.9 | 12.9 ± 1.3 | 25.6 ± 2.5 | 9.5 ± 1.1 |
| Cisplatin + Zingiberofficinale (5 mg/kg + 200 mg/kg) | 30.2 ± 3.2# | 22.1 ± 2.3# | 45.6 ± 4.2† | 15.1 ± 1.7# |

Values are mean presented as Mean ± SEM of six replicate determination. *P < 0.05 significant different when compared to Control group; and ^#^P < 0.05 when compared to Cisplatin induced group

**Table 7.** *The effects of Zingiberofficinale juice extract on cisplatin-induced toxicity on apoptotic markers*

| Group | Caspase-3 (pg/mg) | Bax (pg/mg) | Bcl-2 (pg/mg) |
| --- | --- | --- | --- |
| Control | 0.5 ± 0.1 | 10.2 ± 1.2 | 20.5 ± 2.3 |
| Cisplatin (5 mg/kg) | 1.8 ± 0.3* | 25.6 ± 3.1* | 10.2 ± 1.4* |
| Zingiberofficinale (500 mg/kg) | 0.3 ± 0.1 | 8.5 ± 1.1 | 22.9 ± 2.6 |
| Cisplatin + Zingiberofficinale (5 mg/kg + 200 mg/kg) | 1.2 ± 0.2# | 18.2 ± 2.3# | 15.6 ± 1.9# |
| Cisplatin + Vitamin C (5 mg/kg + 100 mg/kg) | 1.5 ± 0.3# | 22.1 ± 2.8# | 12.9 ± 1.6# |
| Zingiberofficinale + Vitamin C (500 mg/kg + 100 mg/kg) | 0.4 ± 0.1 | 9.5 ± 1.3 | 21.2 ± 2.4 |
| Cisplatin + Zingiber extract (5 mg/kg + 200 mg/kg) | 1.4 ± 0.3# | 20.5 ± 2.6# | 14.2 ± 1.8# |
| Zingiber extract (200 mg/kg) | 0.6 ± 0.2 | 11.9 ± 1.5 | 19.5 ± 2.2 |

Values are presented as mean ± SEM of six replicate determinations. *P < 0.05 significantly different when compared to Control group;and ^#^P <0.05 when compared to Cisplatin-induced group.
